# Ligand Geometry Controls Adhesion formation via Integrin Clustering

**DOI:** 10.1101/435826

**Authors:** Rishita Changede, Haogang Cai, Shalom Wind, Michael P. Sheetz

## Abstract

Integrin-mediated cell matrix adhesions are key to sensing the geometry and rigidity of the extracellular environment to regulate vital cellular processes. *In vivo*, the extracellular matrix (ECM) is composed of a fibrous mesh. To understand the geometry that supports adhesion formation on fibrous substrates, we patterned 10 nm gold-palladium single lines or pairs of lines (total width within 100 nm), mimicking thin single ECM fibers or a minimal mesh geometry, respectively and functionalized it with integrin binding ligand Arg-Gly-Asp (RGD). Single lines showed reduced focal adhesion kinase (FAK) recruitment and did not support cell spreading or formation of focal adhesions, despite the presence of a high density of integrin-binding ligands. Using super resolution microscopy, we observed transient integrin clusters on single lines, whereas stable 110 nm integrin clusters formed on pairs of lines similar to those on continuous substrates. This indicated that two-dimensional ligand geometry is required for adhesion formation on rigid substrates. A mechanism to form modular 100nm integrin clusters bridging the minimal fiber mesh would require unliganded integrins. We observed that integrin mutants unable to bind ligand co-clustered with ligand-bound integrins when present in an active extended conformation. Thus, these results indicate that functional integrin clusters are required to form focal adhesions and unliganded integrins can co-cluster to bridge between thin matrix fibers and can form stable integrin adhesions on dense fibrous networks.

## Introduction

Attachment of cells to the extracellular matrix (ECM) in three dimensional environments is imperative for cell functions that regulate growth, differentiation and diseases ^1, 2^. This attachment is mediated by transmembrane heterodimeric integrin receptors that bind to ligands such as Arginine-Glycine-Aspartate peptides (RGD) on ECM fibrous mesh of Laminin, Collagen or Fibronectin^3^. Most well studied cell ECM attachments are large focal adhesions formed by fibroblasts on two dimensional glass substrates. However, these are rarely observed in *invivo* three dimensional substrates^4^. Architecture of the adhesions observed *invivo* are small and punctate^5^. This could be due to the fibrous architecture of the 3D *in vivo* ECM that could be composed of small fibers 2-20 nm in diameter ^6, 7^. However, the formation of cell matrix adhesions on these substrates is poorly understood. In particular what is the role of ligand density versus ligand geometry in adhesion formation? Studies on minimal ligand density needed for cell spreading revealed that clusters of 4-7 RGD functionalized 5 nm nanodots spaced within 100 nm supported cell spreading^8, 9^, except on very soft substrates^10^, possibly through the assembly of clusters of adhesion proteins^11, 12^. Further, in context of fibrous mesh ECM, what is the minimal fibrous ligand geometry that would support adhesion formation?

The mechanism of how the minimal dot geometry supports adhesion formation is not understood. In minimal density experiments, few integrin ligands could support formation of adhesions. To understand the molecular basis of this, we hypothesized that co-clustering of a few ligand-bound integrins with unliganded integrins could increase the avidity of ligand-integrin binding as well as sensitize the cell to low ligand densities through a threshold-based signaling complex^13–15^. Several previous findings supported this hypothesis. A significant fraction of integrins were motile within the focal adhesions, suggesting that non-ligand bound integrins were present within the focal adhesion^16^. Further, it has been shown that chimeric receptors with integrin cytoplasmic tails and extracellular and transmembrane domains of other receptors (either cadherin of interleukin-2 receptor alpha) localized to the focal adhesions indicating that all integrins in adhesions were not necessarily ligand-bound ^17, 18^. Within mature adhesions, a few integrins exerted high forces, whereas a significant fraction of them were in the low force state indicating that a few ligand-bound integrins were sufficient for adhesion formation^19^. Those mature adhesions typically formed from modular nascent adhesions after rigidity-sensing^20^.

Cell matrix adhesion formation is initiated by formation of nascent adhesions ^21, 22^ of ~100 nm in diameter ^23–25^ that activated differential mechanotransduction signals based on substrate rigidity^26, 27^. To address the minimal geometry needed for adhesion formation in fibrous ECM, we patterned glass substrates with narrow lines of gold palladium (AuPd) mimicking one dimensional individual 10 or 20 nm single fibers or a pair of lines spaced within the diameter of a single modular integrin cluster of ~100 nm ^24^ mimicking a minimal two dimensional mesh within it. We tested if cell matrix adhesions can form on single 10 or 20 nm lines. Unlike complex 3D matrices, fabricated substrates change a single parameter at a time allowing us to probe the geometric organization of ECM fibers required for formation of cell matrix adhesions.

Cells did not spread if RGD functionalized dots were 100 nm or more apart. Thus, formation of modular adhesion clusters of ~100 nm appeared to be an important aspect of robust adhesion formation^23^. Within the clusters many weak integrin-ligand bonds were integrated to form a strong adhesive patch that supported a range of force densities in response to different mechanical properties of the matrix ^28^. Hence we tested if formation of modular integrin clusters was critical for formation of mature cell matrix adhesions. We used PhotoActivated Localization Microscopy (PALM) to determine if modular integrin clusters could form on fiber-mimic substrates. The dynamics of these adhesion clusters were addressed using structured illumination confocal microscopy (confocal iSIM, see methods).

Having established the geometric requirements (2D vs. 1D) for cluster formation, we probed the mechanisms of cluster formation. It was not known if and how unliganded integrins could aid in the assembly of modular integrin clusters. In order to test this hypothesis, we used point mutants in the β_3_ integrin that were either constitutively activated (N305T)^29^ or could not bind ligand (D119Y)^30^ and tested whether ligand binding or integrin activation associated with integrin clusters to aid their formation. Hence the goal of this study was to understand the geometric requirement of the ECM to form cell matrix adhesions and the mechanism of how small adhesions could bridge the narrow ECM fibers.

## Results

### Adhesion clusters require two dimensional ligand geometry to assemble and stabilize

To determine whether single matrix fibers or fiber networks were needed for adhesion formation, single 10 nm AuPd lines or pairs of lines separated by 80 nm were patterned by electron beam lithography (Figure 1a, 1b). The center-to-center distance between consecutive single lines or line pairs was 500 nm (Figure 1b), approximately five times larger than a single integrin cluster. Based upon previous studies, RGD functionalized hexagonal nanodot patterns enabled cell spreading^8, 9^. Therefore hexagonal nanodot pattern comprising of seven 10 nm dots separated by 40 nm served as a positive control. Individual dot patterns were spaced by 402 nm (nanodot global density of 50/μm^2^, Figure 1a, 1b, see Methods for details). The total line pair width equaled the diameter of the hexagonal dot pattern, which was about the size of a single modular integrin cluster (~100 nm). AuPd was selectively functionalized with cyclic RGD using thiol and biotin/ neutravidin chemistry (Figure 1c)^31^. The integrated ligand density over 25 μm^2^ for single lines was higher than the density for hexagonal dot patterns, and the density of line pairs was about twice that of single lines (Figure 1d, S1a, S1b). The glass substrate was passivated using 1,2-dioleoyl-sn-glycero-3-phosphocholine (DOPC) supported lipid bilayers to prevent non-specific ligand binding^32^. To assay formation of cell matrix adhesions, either human foreskin fibroblasts (HFFs) or mouse embryonic fibroblasts (MEFs) were plated on the patterned substrates and allowed to spread for 30 minutes (Supplementary Figure S2a). Similar results were obtained using both cell lines, demonstrating that the interactions were not specific for a given cell line of fibroblasts.

**Figure 1:**
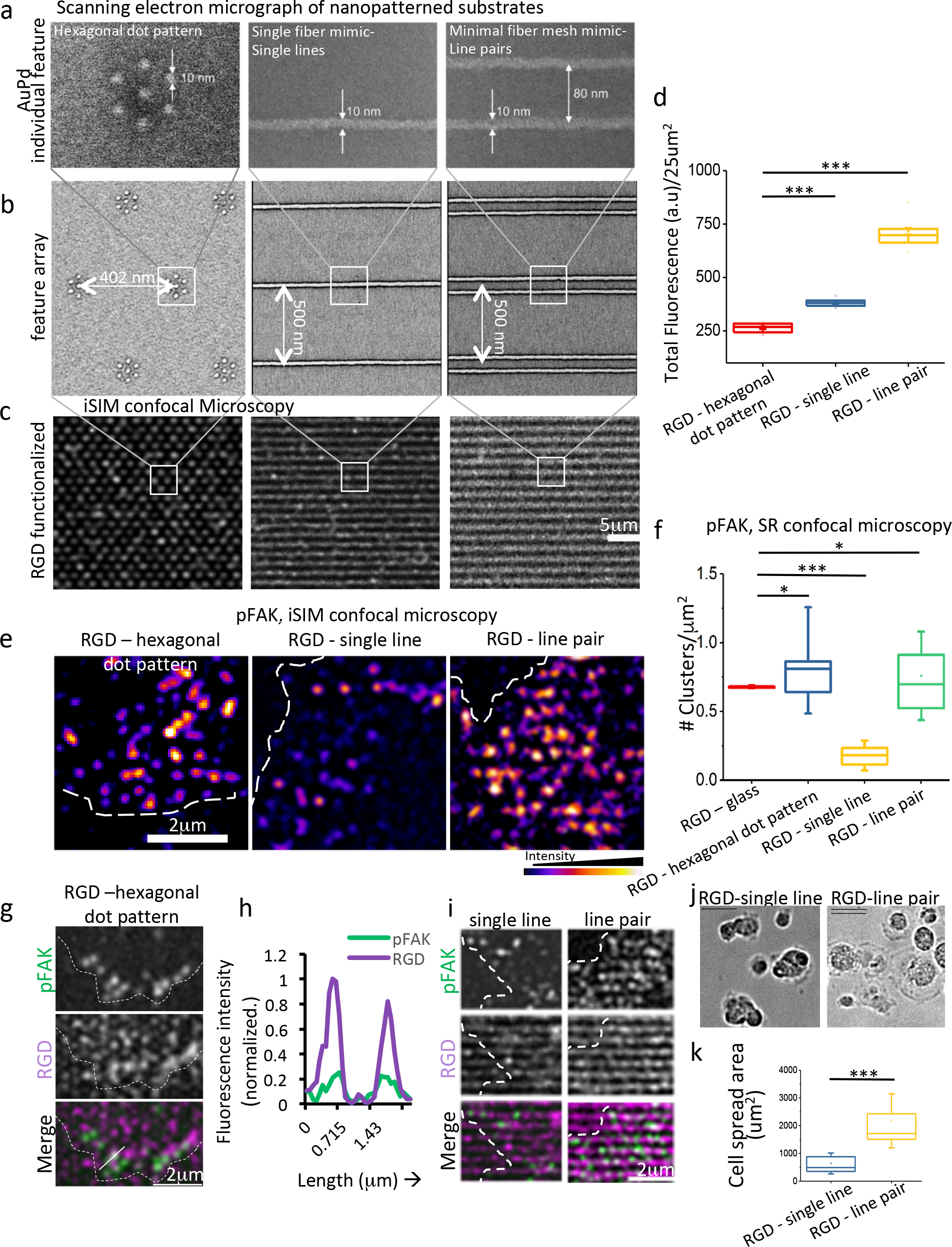
Focal adhesion formation and cell spreading was reduced on nanopatterned substrates mimicking single thin matrix fibers. (a) scanning electron microghaph shows a single feature of AuPd patterned substrates - hexagonal dot pattern (left panel), single fiber mimetic single 10 nm line (middle panel) and minimal fiber mesh mimetic line pair (b) shows false color array of features before lift-off during the patterning process. White box shows the zoom in region depicted in the panel above (c) super resolution confocal image of RGD functionalized substrates. (d) Integrated fluorescence intensity (arbitrary units) over 25um^2^. (e) False color image of human foreskin fibroblasts (HFF) spread on RGD functionalized nanopatterns for 30 minutes and immunolabeled with pFAK397. Dotted while line marks the cell edge. (f) shows the number of immunolabeled phosphoFAK clusters/um2 of the lamellipodial region as marked by 4um from the cell edge when cells spread on the various RGD functionalized substrates, n>7cells (g & i) top panel shows phosphoFAK (397) immunolabeling in green, middle panel shows RGD in magenta and the bottom panel shows the merge (h) normalized fluorescence intensity of RGD and phosphoFAK through the line depicted in Merge. (j) Top panel shows HFFs spread on single lines (left panel) or line pairs (right panel) and immunolabeled with phosphoFAK. In all panels line at the top left corner represents the orientation of the fiber mimetics, dotted line marks the cell edge. (k) Cell spread area on fiber mimetics, n>10cells. Box plots showed upper bound at 75% and lower bound at 25% (box), ±SD (whiskers). Line marked the as median and dot marked the mean of the population. The indicated p values were obtained using two-tailed Student’s t-tests. *p*>*0.5 n.s*, *0.5*>*p*>*0.1*\*, *0.1*>*p*>*0.001*\*\*, *p*<*0.001*\*\*\*.

In HFFs, immunolabeling revealed discrete phosphoFAK (Y397) clusters (Figure 1e, 1f). They were imaged using SR confocal microscopy, which increased the resolution of confocal microscopy by two-fold and enabled us to resolve individual adhesion clusters^24^. On line pairs and hexagonal dot patterns, cluster density in the lamellipodium was comparable to those for cells spread on continuous glass substrates. Significantly fewer clusters were observed on the single line patterns (Figure 1e, 1f, S4a), indicating that activated phosphoFAK signaling was significantly reduced and focal adhesions did not form.

Overlap of phosphoFAK and RGD (Figure 1g, 1h) showed that functional adhesions formed over the RGD-hexagonal dot patterns despite the low density of RGD ligands when the ligands were distributed within ~100 nm in two dimensions. Similar clusters of phosophoFAK were observed in adhesions formed on continuous RGD-coated glass substrates (Supplementary Figure S3a). Fewer clusters were observed on single lines but the clusters were formed overlapping with the nanopatterned lines (Figure 1i). We also noted orthogonal alignment of focal adhesion with respect to the orientation of the line pairs, emphasizing that a spacing of 500 nm did not hinder the formation of mature adhesions (Supplementary Figure S3b). Cell spreading was poor on single fiber mimetic substrates (Figure 1j, 1k; spreading area - 640 ± 360 μm^2^ on single lines, compared to 2175 ± 900 on line pairs), and fewer overall adhesion were formed (Supplementary Figure S2b). As a positive control, neighbouring cells bound to a continuous AuPd patches separating the single line patterns and formed larger adhesions (Supplementary Figure S2b). On 20 nm single lines and line pairs, similar results compared to 10 nm lines were obtained respectively (Supplementary Figure S4a, S4b). This indicated that high ligand density presented in one dimension was not sufficient and local ligand geometry played a significant role in formation of cell matrix adhesions.

To validate the effect of local ligand geometry, single lines were spaced by 250 nm (~2.5 times larger than an integrin cluster) and line pairs were spaced by 500 nm to give a comparable effective concentration of ligand at the length scale of a single cell (Supplementary Figure S5a, S5b, S5c). Again, MEFs spread poorly on single lines and formed smaller adhesions compared to line pairs (Supplementary Figure S5d). To compensate for possible lower integrin activation on single lines, we treated the MEFs with manganese chloride (MnCl_2_), a potent activator of α_v_β_3_ integrin^33^, and spread them on single lines versus line pairs. This was not sufficient to recover the cluster formation or adhesion formation on single lines (Supplementary Figure S6a, S6b). This showed that local ligand geometry dictated cell adhesion formation even if overall ligand density was constant. One-dimensional geometry mimicking thin, single ECM fibers was not sufficient to support adhesions.

Reduction of focal adhesions and FAK signaling on single lines could be related to reduction in size of modular integrin clusters. We assayed integrin clusters using PALM microscopy. MEFs expressing β_3_-PAGFP were spread on nanopatterned substrates for 30 minutes and fixed. Continuous RGD functionalized glass and polylysine-coated glass served as positive and negative controls respectively. Similar clusters appeared on hexagonal dot patterns (FWHM was 114±11 nm, mean±SD) and continuous RGD (108±12 nm, Figure 2a, 2b). In contrast, less dense, smaller clusters appeared on polylysine coated glass (73±17; Figure 2a, 2b). This was also evident in the intensity of PAGFP in a clustered region as a function of #frames in the PALM image. Multiple peaks were seen in the case of hexagonal dot patterns and RGD coated glass, as opposed to polylysine-coated substrates (Supplementary Figure S7a, S7b) showing that integrin clusters formed within focal adhesions.

**Figure 2:**
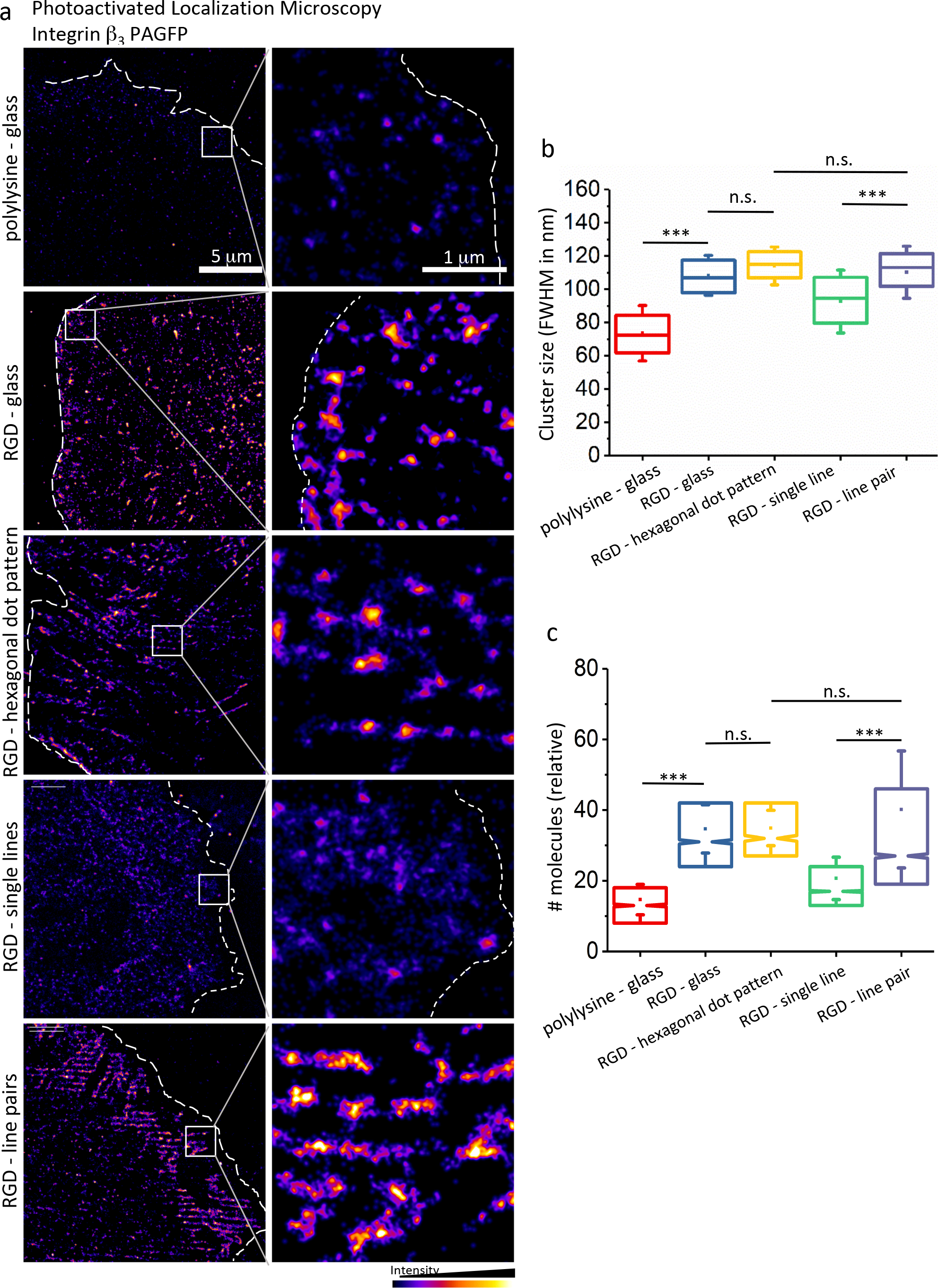
Reduction in phosphoFAK signaling was correlated with fewer integrin clusters on the single matrix fiber mimetics. (a) Photoactivated light microscopy (PALM) image of MEFs expressing β_3_-PAGFP spread on various substrates. Left panel shows MEFs, dotted line marks the cell boundary and white box marks the zoom in region shown in the right panel. Line of the top left represent the orientation of the nanopatterns. n>4 cells (b) size of clusters (full width half maxima, nm) on the various substrates and (c) relative number of molecules within the cluster on various substrates. Number of clusters >800 from 4 or more cells. Box plots showed upper bound at 75% and lower bound at 25% (box), ±SEM (whiskers). Line marked the as median and dot marked the mean of the population. The indicated p values were obtained using two-tailed z test. *p*>*0.5 n.s*, *0.5*>*p*>*0.1*\*, *0.1*>*p*>*0.001*\*\*, *p*<*0.001*\*\*\*.

On the line patterns, linear arrays of integrin clusters formed within adhesive regions. On the single lines, fewer integrin clusters were observed compared to line pairs (Figure 2a) and the size of the clusters was smaller (93±18 nm) compared to clusters on the line pairs (112 ±14 nm, Figure 2b). Although the absolute number of integrins in the clusters was difficult to determine due to the presence of endogenous unlabeled integrins, incomplete folding of PAGFP and stochastic blinking errors^34^, it was possible to compare the relative numbers of photoactivated integrins within the clusters. On the hexagonal dot pattern and on the continuous RGD-coated glass, clusters contained a significantly higher number of integrins (35±11 mean ±SD) than on polylysine coated glass (mean of 15±9) (Figure 2c). On line pairs, the number of molecules in the clusters (40 ± 23) was similar to that on hexagonal clusters and the continuous RGD glass (Figure 2a, 2c) whereas there were significantly fewer molecules in clusters on single lines (24±23). Thus, even though the density of ligands with 10 nm lines was greater than with hexagonal dot patterns, the clusters were significantly smaller and had fewer integrins. This raised the question of whether adhesions on single lines were transient or did not form at all.

### Adhesion clusters are transient on single lines

To determine if adhesion assembly, lifetime or both were decreased on single lines, we tracked the paxillin-mApple distribution in cells spread on both line geometries. Paxillin localized to adhesion clusters on both single lines and line pairs, but fewer clusters formed on the single lines (Figure 3a; Supplementary Movie S1) as compared to line pairs (Figure 3d, Supplementary Movie S2). Time lapse imaging also revealed that clusters on single lines were transient (Figure 3b, 3c) and disassembled rapidly, with a lifetime of about ~100 seconds from the kymographs (Figure 3c, 3k). On line pairs, most clusters were long lived, with an average lifetime greater than 480 seconds, the total time of imaging (Figure 3e, 3k). Thus, more adhesions formed on double lines and they lasted for a much longer time than on the single lines.

**Figure 3:**
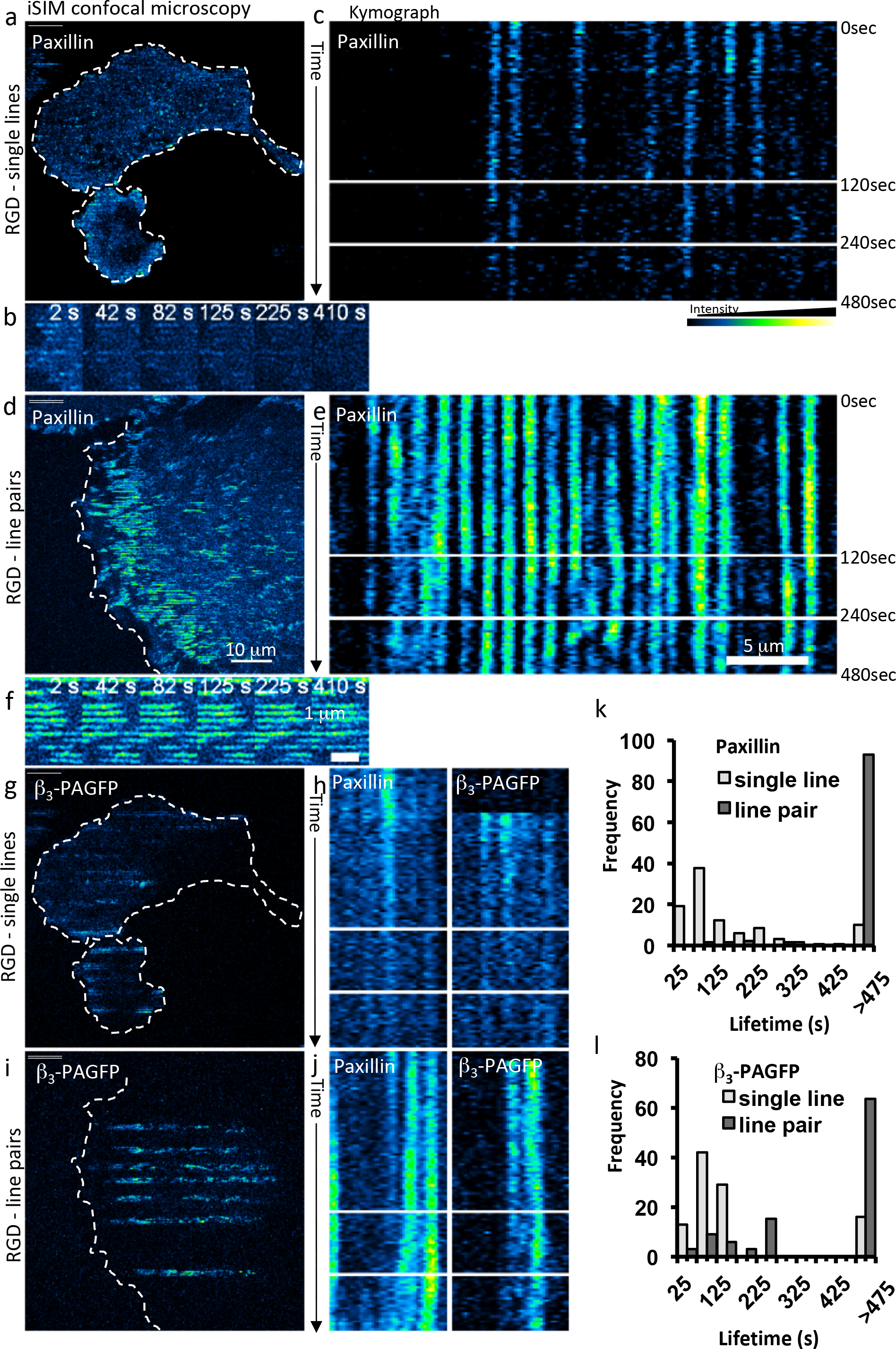
Stable adhesion clusters were formed on line pairs but not on single lines. (a, d) MEFs spread on single lines or lines pairs imaged in real time using Paxillin mApple as a label for adhesions. (b, e) zoom in region over time of the rectangle marked in and e (c, f) Kymograph of a line across adhesion clusters depicted in b, f respectively. From top to first line, each pixel represents 1 second interval for 120 seconds, from first black line to second black line, its 5 second interval for 120seconds and from the second black line to the bottom time interval is 10seconds for 240 seconds, total length of the movie is 480 seconds (g, i) Same cell as shown in a and d, where β_3_-PAGFP was activated in several horizontal region of interest. (h, j) Kymographs of β_3_-PAGFP in regions marked as straight line, time similar to c. (k, l) frequency plots of lifetime of clusters marked with mApple Paxillin or b3-PAGFP respectively, on single lines (light grey) or line pairs (dark grey) n>5 cells and >80 clusters.

To further address specific integrin clustering, we used β_3_-PAGFP and photoactivated it in linear regions of interest parallel to the substrate lines (Figure 3i-l). Fluorescence Loss After photoactivation (FLAP) revealed that on single lines, fewer integrins were observed in the adhesion regions (Figure 3g) and these integrins left clusters rapidly, as observed by the kymograph (Figure 3h). This measurement also showed that clusters had an average lifetime of ~100 seconds (Figure 3l), although a few were long-lived. In contrast, on the line pairs, integrin clusters had many more integrins (seen by higher intensity) and were long-lived (Figure 3i, 3j) with average lifetime greater than 480 seconds (Figure 3l).

### Unliganded Integrins Assemble in Clusters with Ligand bound Integrins

Formation of stable adhesions clusters was essential for adhesion formation and clusters of comparable size formed on hexagonal dot patterns and line pairs. Integrin extracellular domain in the extended confirmation is ~20 nm, significantly shorter than the distance between the nanopatterns. Yet, dense integrin clusters appear to bridge the nanodots or line pairs indicating that unliganded integrins could assemble within these clusters to reach the modular size. To test this hypothesis, we used RGD functionalized supported lipid bilayers (see methods), since adhesion clusters were previously observed to be long-lived on them and it allowed us to image using two color PALM ^24, 35^. We used integrin point mutant β_3-_D119Y that did not bind RGD ligand (Figure 4a) and the mutant β_3_N305T that was in a constitutively activated form both labeled with mEOS2. With two color PALM, we asked if mutant integrins (labeled with mEOS2) localized with endogenous integrins clusters (observed by clustering of RGD ligand).

**Figure 4:**
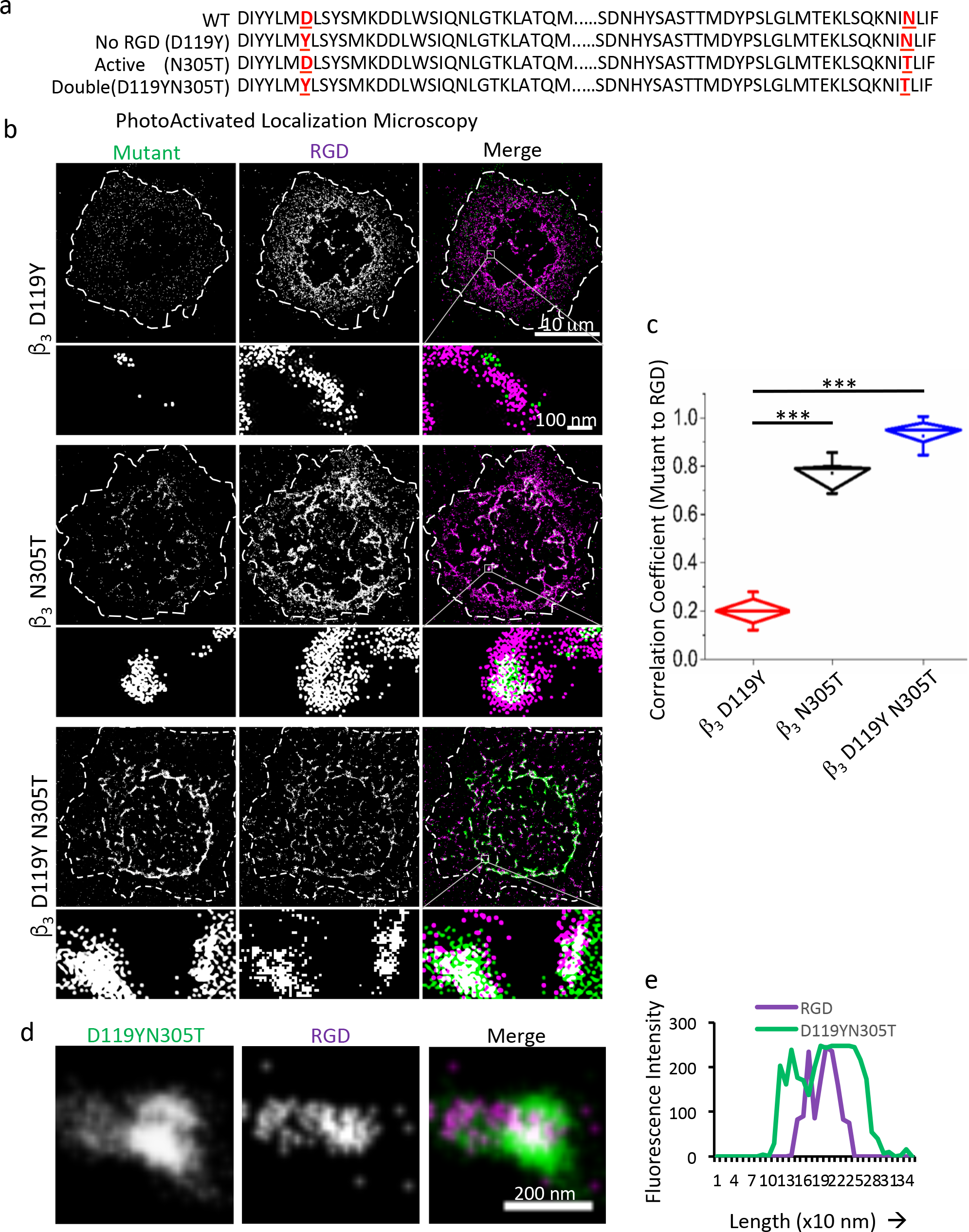
Activation is sufficient and ligand binding is not necessary for integrins to localize to adhesion clusters. (a) mutants of β_3_ integrin in the extracellular domain, D119Y which cant bind RGD ligand, N305T constitutively active and double mutant D119YN305T. (b) MEF cells expressing mutant β_3_ and endogenous β_3_ spread on RGD functionalized supported lipid bilayer. Two color PALM image of the mutant β_3_ mEOS2 and RGD labeling the endogenous integrin clusters. Dotted line marks the cell edge and the white box marks the region zoomed shown in the panel below. (c) Correlation coefficient of β_3_ mutant and endogenous β_3_-RGD clusters n>15cells. (d) zoom in region of a clustered region of D119YN305T(e) intensity plot of the line ROI draw in d, merge panel. Diamond plots showed upper bound at 75% and lower bound at 25% (diamond), ±SD (whiskers). Line marked the as median and dot marked the mean of the population. The indicated p values were obtained using two-tailed Student’s t-tests. *p*>*0.5 n.s*, *0.5*>*p*>*0.1*\*, *0.1*>*p*>*0.001*\*\*, *p*<*0.001*\*\*\*.

In MEFs, we overexpressed mutant β_3_ isoforms without altering the endogenous β_3_ and spread the cells on RGD functionalized lipid bilayers for fifteen minutes. The non-ligand-binding integrin did not colocalize with the endogenous integrin (Figure 4b, Pearson’s correlation coefficient-0.2 Figure 4c) possibly because it was not activated. The constitutively activated integrin interspersed with the ligand clusters since it bound ligand (Figure 4b) and showed a high degree of colocalization (0.7, Figure 4c). (This also confirmed that the fluorescent tag did not cause an exclusion from the adhesion clusters.) In order to test whether the failure of the D119Y mutant to colocalize with the clusters was related to its conformation (i.e., that it was in a bent, inactive conformation due to the absence of ligand stabilizing the active confirmation), we produced a doubly mutated integrin that was unable to bind ligand but was locked in an activated form (β_3_ D119Y N305T-mEOS2). This double mutant colocalized with the endogenous adhesion clusters (Figure 4a, 4b, 4c) with a very high Pearson’s correlation coefficient (mean 0.8), comparable to β_3_N305T. Further, not only did it colocalize with the adhesion clusters, but it was often present around the RGD, implying that unliganded integrins were recruited to the clusters formed by the ligand bound integrins (Figure 4d, 4e). Thus, activated but unliganded integrins were able to associate with modular adhesion clusters, providing a mechanism for the formation of adhesion clusters on the sparsely presented ligands of line pairs and dot arrays.

## Discussion

The results reported here demonstrate that isolated linear ligand arrays (10 or 20 nm in width) are unable to support matrix adhesions even if the density of ligands is high enough. When few RGD ligands^3^ are in hexagonal dot pattern within about 100 nm, they support the formation of strong adhesion clusters of comparable size to those observed on continuous RGD coated glass. In addition, when two 10 nm lines are brought within the diameter of a single cluster, they catalyze stable adhesion assembly similar to RGD-coated glass. Super-resolution microscopy indicates that unliganded integrin molecules assemble in adhesion clusters and bridge between ligand bound integrins on adjacent lines or dots. These different RGD patterns show that ligand geometry is an important factor in modular adhesion cluster formation and subsequent adhesion development on rigid substrates: only a few ligands are needed to form a stable cluster, but a two-dimensional distribution is necessary (Figure 5). This indicates that the single small matrix fibers (≤ 20 nm) are unable to support adhesion formation as they destabilize the modular adhesion clusters. The modular nature of the adhesion proteins (FAK, β_3_ integrin) is retained in adhesions on varying geometries suggesting that it forms the fundamental basis of cell matrix adhesions and could be required for signaling^23^. In adhesions, the closely spaced but discrete integrin clusters could catalyze formation of multiple adhesion protein complexes by producing high local concentrations of signaling components while still enabling free access of inactivating proteins and allowing for a large and varied number of protein-protein interactions needed for signaling via the adhesion ^36–39^.

**Figure 5:**
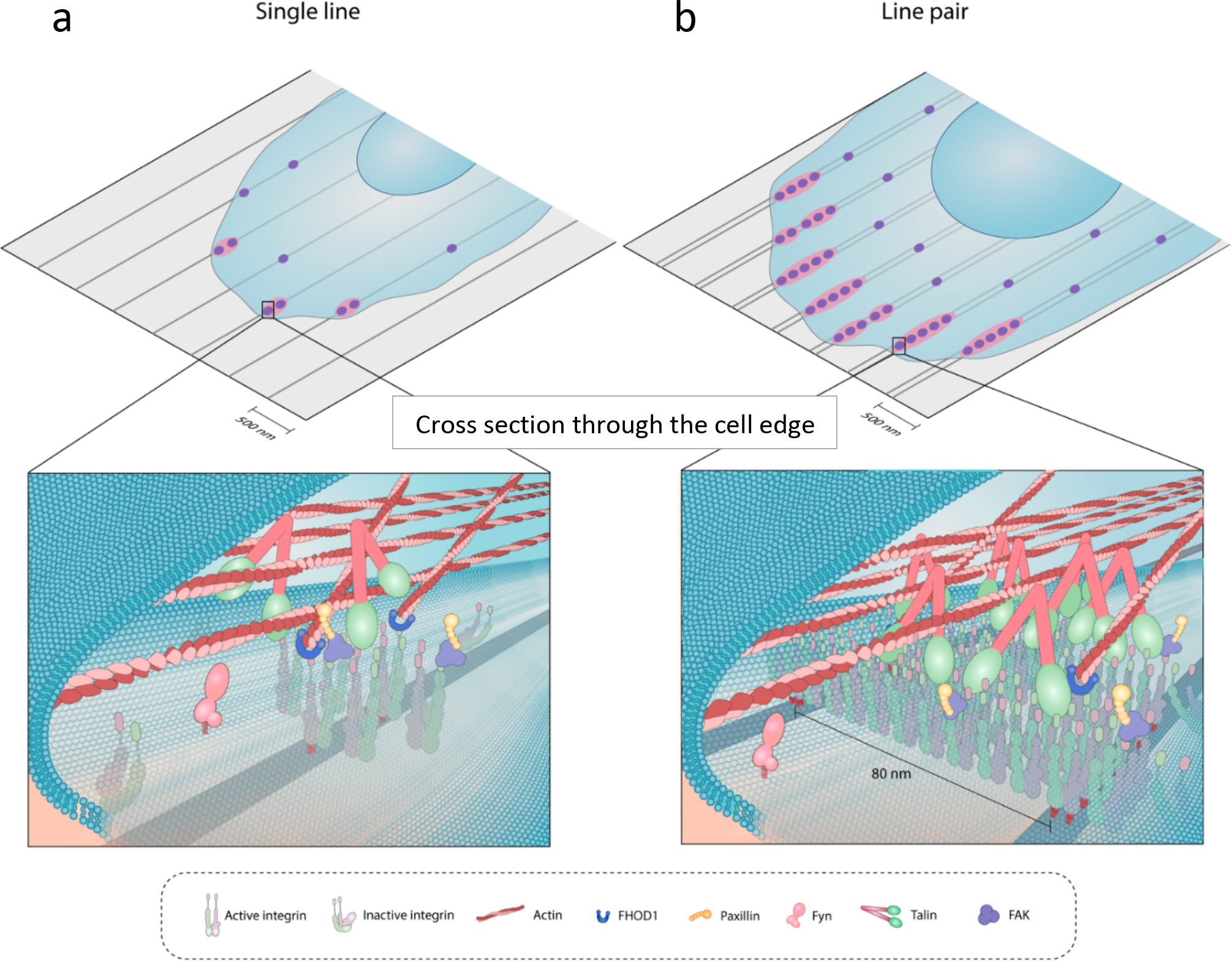
Proposed model for assembly of adhesion clusters. (a) On single lines, even in presence of ligands, few transient adhesion clusters assemble that cannot support cell spreading, suggesting single ECM fibers are not sufficient for cell spreading. (b) On line pairs stable uniform adhesion clusters assemble that are composed of ligand bound and unliganded integrins. These adhesion clusters assemble into larger focal adhesions, implying that integrin clustering and two dimensional ligand geometry are required for formation of focal adhesions on rigid substrates.

From the initial stages of cluster formation, there is a rapid sequence of steps that will determine if the nascent adhesion matures into a focal adhesion, a podosome, or will disassemble ^40^. The very transient nature of the integrins on the single lines indicate that they are not normal clusters. This is surprising since the density of RGD was greater on the single lines than on the hexagonal dot patterns (Figure 2c). One possible explanation for the low level of adhesion formation and the rapid turnover is that the linear arrays are unable to support forces on the activated integrin molecules in forming clusters. Thus, we suggest that the on rigid substrates single lines do not form sufficiently stable adhesions to enable cells to develop forces on adhesion clusters that consequently dissipate (Figure 5).

There are two major hypotheses to explain why clusters are limited to 100 nm in diameter over a wide range of ligand densities and changes in talin and actin at the cell matrix adhesion^24^. First, there is a two-dimensional cytoplasmic adhesion protein complex that stabilizes the integrins in the cluster. From measurements of talin length in cells, the molecules are typically in a trigonal configuration spanning over 100 nm (Figure 5) ^41^, and talin mutations can change the cluster diameter ^24^. The second hypothesis claims that the integrin clusters bend the membrane until a limiting curvature is reached, since integrins partition preferentially to curved membrane regions ^42^. In both of these models, there are reasonable explanations for why cluster stability would be greatly increased by a two-dimensional arrangement of ligands. These hypotheses are not mutually exclusive, and formation of modular adhesion clusters could be a resultant of several processes activated simultaneously.

Most matrix proteins assemble into fibers that can range in diameter from 2 nm to several micrometers for large collagen fibers. Based upon our findings, there must be dense arrays of parallel small fibers such that the total width is within ~ 100 nm in some regions. However, overlapping fiber arrays should support adhesions at sites of fiber overlap. One-dimensional stress on fiber arrays should increase junctions and adhesion contacts. On two-dimensional surfaces like basement membranes, a critical local density of ligands is needed and that can be as few as four ligands in a ≤ 60 nm quadrilateral pattern^9^. Such modular adhesions based on single integrin clusters could be the basis of small adhesions observed in three dimensional environments when fibers are overlapping. Since only about ten percent of the integrins need to be ligand bound for adhesion maintenance, the forces on individual small fibers may be very high, and that could aid in matrix remodeling. Fibrosis or excess deposition of large collagen fibers would decrease the force per fiber and compromise the cell’s ability to remodel the collagen. The detailed aspects of two-dimensional integrin clusters logically influence many cell-matrix interactions and those interactions are critical for determining cell viability and function.

## Acknowledgements

We thank Gregory Giannone, Neurosciences Bordeaux, France, and Michael W. Davidson group, The Florida State University, Tallahassee, FL, USA for DNA constructs. We thank Haguy Wolfenson for his help with initial experiments. This work is supported by intramural funds from the Mechanobiology Institute. RC is supported by Singapore National Research Foundation’s CRP grant (NRF2012NRF-CRP001-084).

## Author Contributions

RC and MS conceived and designed the experiments. HC & SW prepared AuPd nanopatterned substrates. RC and HC standardized the functionalization of AuPd substrates with RGD. RC performed the experiments, analyzed the data and wrote the manuscript. R.C., M.P.S., S.W., H.C., prepared the manuscript.

